# Actin Y53 Phosphorylation Regulates Somatic F-actin Radial Translocation to Promote Neuronal Polarization

**DOI:** 10.1101/2023.06.17.545417

**Authors:** Shuai Hong, Durga Praveen Meka, Oliver Kobler, Robin Scharrenberg, Carina Meta Friedrich, Birgit Schwanke, Melanie Richter, Froylan Calderon de Anda

**Author notes:** ^#^ Equal contribution (Shuai Hong, Durga Praveen Meka). To whom correspondence should be addressed: Durga Praveen Meka, Froylan Calderon de Anda). ^$^Lead Contact Froylan Calderon de Anda).

## Abstract

How neurons accomplish the growth of an axon, and several dendrites sequentially is a cellular process not well understood. Here, we show that preferential somatic F-actin delivery to neurites inhibits neurite growth during axon formation. Thus, the neurite receiving less somatic F-actin is growing as an axon. During dendrites outgrowth, radial somatic F-actin translocation into all neurites is drastically diminished suggesting that cellular growth is restricted by the somatic F-actin flow into the growing neurites. Mechanistically, we report that radial somatic F-actin translocation is mediated via the Myosin II motor. Pharmacological inhibition of these molecular motors promotes neurite elongation and precludes F-actin translocation into neurites. Moreover, we show that actin phosphorylation at Y53, which promotes F-actin instability, is enriched selectively in some neurite shafts to exclude/minimize translocation of somatic F-actin only into those neurites. Therefore, enrichment of actin pY53 in the neurite shaft correlates with the longer neurite during axon formation. Accordingly, the Kinesin-1 motor domain, which accumulates transiently in neurites of stage 2 neurons and only in the emerging axon of stage 3 neurons, localizes preferentially in the neurite with the increased actin pY53 signal in the neurite shaft. Finally, we demonstrate that microtubule acetylation promotes actin phosphorylation, and these cytoskeleton modifications coexist in the longest neurite that grows as an axon. Collectively, our data support a model in which somatic F-actin translocation into undifferentiated neurites impedes growth counteracted by actin pY53, consequently, supporting neuronal polarization during axon formation.

## Introduction

Differentiating pyramidal neurons follow an intricate process during which one of the neurites is specified as the axon and the remaining typically become dendrites. This cellular process is highly stereotyped in cultured neurons and *in situ* with the axon growing first followed by the dendrite’s development ^1–3^. One important question in neurobiology is how early developing neurons cope to sustain the elongation of only one neurite during axon formation. In other words, why do dendrites and axon not grow simultaneously? It was suggested an intrinsic feedback system with inhibitory internal signals precludes neurite extension of future dendrites during axon elongation. On the other hand, positive molecular cascades might sustain locally the axon extension ^4^. Thus, a local increase of cAMP in one neurite decreases cAMP in all other neurites of the same neuron. The alterations in the cAMP and cGMP levels in the neurites are mutually opposing. Consequently, local, and long-range reciprocal regulation of cAMP and cGMP together ensure coordinated development of one axon and multiple dendrites ^5^. Whereas semaphorin 3A acts as an inhibitory signal that suppresses axonal growth (and thus promotes dendritogenesis) in cultured hippocampal neurons ^6^. Long-range inhibitory signaling mediated by Ca^2+^ waves suppress the outgrowth of minor processes by activating RhoA thereby ensuring neuronal polarization ^7^. These data suggest that positive feedback signals are continuously activated in one of the minor neurites resulting in axon specification and elongation. In turn, negative feedback signals are propagated from a nascent axon terminal to all minor neurites to inhibit their outgrowth at the same time, thereby leading to asymmetric growth. All the downstream signaling mechanisms suggested for the positive and negative feedback signals seem to converge at two fundamental aspects related to cytoskeleton dynamics. 1. Increased microtubule stabilization and 2. Enhanced F-actin dynamics (actin instability) ^4, 8, 9^.

Recently, we showed that somatic F-actin which concentrates near the centrosome in a form of F-actin asters is delivered to the growing neurites following an inversely proportional correlation to the length of the neurites ^10^. Hence, suggesting a somatic F-actin translocation into growth cones that could act as a growth-inhibiting factor. Along these lines, the removal of F-actin from growth cones/neurite tips by pharmacological means promotes neurite outgrowth ^11, 12^. Based on these observations, we now hypothesize that the somatic organization of F-actin preferentially provides F-actin to peripheral places where the outgrowth should be limited (eventually the shorter neurites), thereby acting as an inhibitory or a pulling force, which suppresses neurite growth. On the other hand, places receiving less F-actin, but more stable microtubules ^13–15^, should be primed for growth, leading to asymmetrical outgrowth/axon formation.

Mechanistically, we showed that this preferential delivery of somatic F-actin into shorter neurites during axon formation is dependent on Myosin II. Consequently, pharmacological inhibition of this motor promotes neurite elongation. Furthermore, we described that actin phosphorylation at Y53, which promotes F-actin instability ^16, 17^, in the neurite shaft precludes somatic F-actin translocation into growth cones. Thus, we detected enriched actin phosphorylation at Y53 in the growing axon which received less somatic F-actin. Finally, we found that microtubule acetylation, which is enriched in the growing axon ^15, 18^, was coupled with actin phosphorylation at Y53. Hence, pharmacological enhancement of microtubule acetylation increased actin phosphorylation at Y53. Altogether, our data provide a new model that explains the neurite growth restriction of the future dendrites during axon elongation.

## Results

### Somatic F-actin translocate to neurites to restrict growth

Axon formation is a hallmark of neuronal polarization in early developing hippocampal and cortical pyramidal neurons ^1–3, 19, 20^. Neurons immediately after plating are round forming lamellipodia (stage 1), then neurons become multipolar extending several neurites (stage 2; ^1^), a morphology which is also found in the developing cortex ^2, 3, 20, 21^, from which usually the one neurite with the fastest growth rate becomes the axon (stage 3; ^1^), while the remaining neurites transform into dendrites (stage 4; ^1, 22^). It is not well understood, however, how neurons cope to elongate first the axon and then the dendrites. We now test the hypothesis that somatic F-actin serves as a negative signal for neurite growth. For that aim, we used stage 2-4 cultured neurons transfected with photoactivatable GFP-Utrophin (PaGFP-UtrCH), which specifically labels F-actin ^23, 24^. PaGFP-UtrCH photoactivation is irreversible in response to 405 nm light with an emission peak at 517 nm. We followed the same protocols for photoactivation in the cell body previously reported by us ^10^ and we found that in stages 2 and 3 neurons photoactivated PaGFP-UtrCH segregated preferentially into minor neurites but not the growing axon (Figure 1a-b; Videos 1-2). At stage 4, however, when dendrite outgrowth was undergoing, the somatic photoactivated PaGFP-UtrCH signal remained in the cell body without detection of PaGFP-UtrCH photoactivated signal in neurite shafts (Figure 1c, d; Video 3). These results show that the somatic photoactivated PaGFP-UtrCH signal translocated into neurites is following an inverse correlation with growth. Consequently, we decided to test whether somatic F-actin translocation into neurites inhibits growth. To this end, we performed somatic PaGFP-UtrCH photoactivation in stage 2-3 neurons to track the photoactivated PaGFP-UtrCH signal translocation into neurites and monitor neurite growth afterwards (1-3 hrs.). We found that neurites receiving less photoactivated PaGFP-UtrCH signal grew after 1-3 hrs. after photoactivation (Figure 2a-f). On the contrary, neurites receiving more photoactivated PaGFP-UtrCH signal retracted (Figure 2a-f). These results strongly suggest that somatic F-actin could serve as an inhibitory signal for neurite growth.

**Figure 1.**
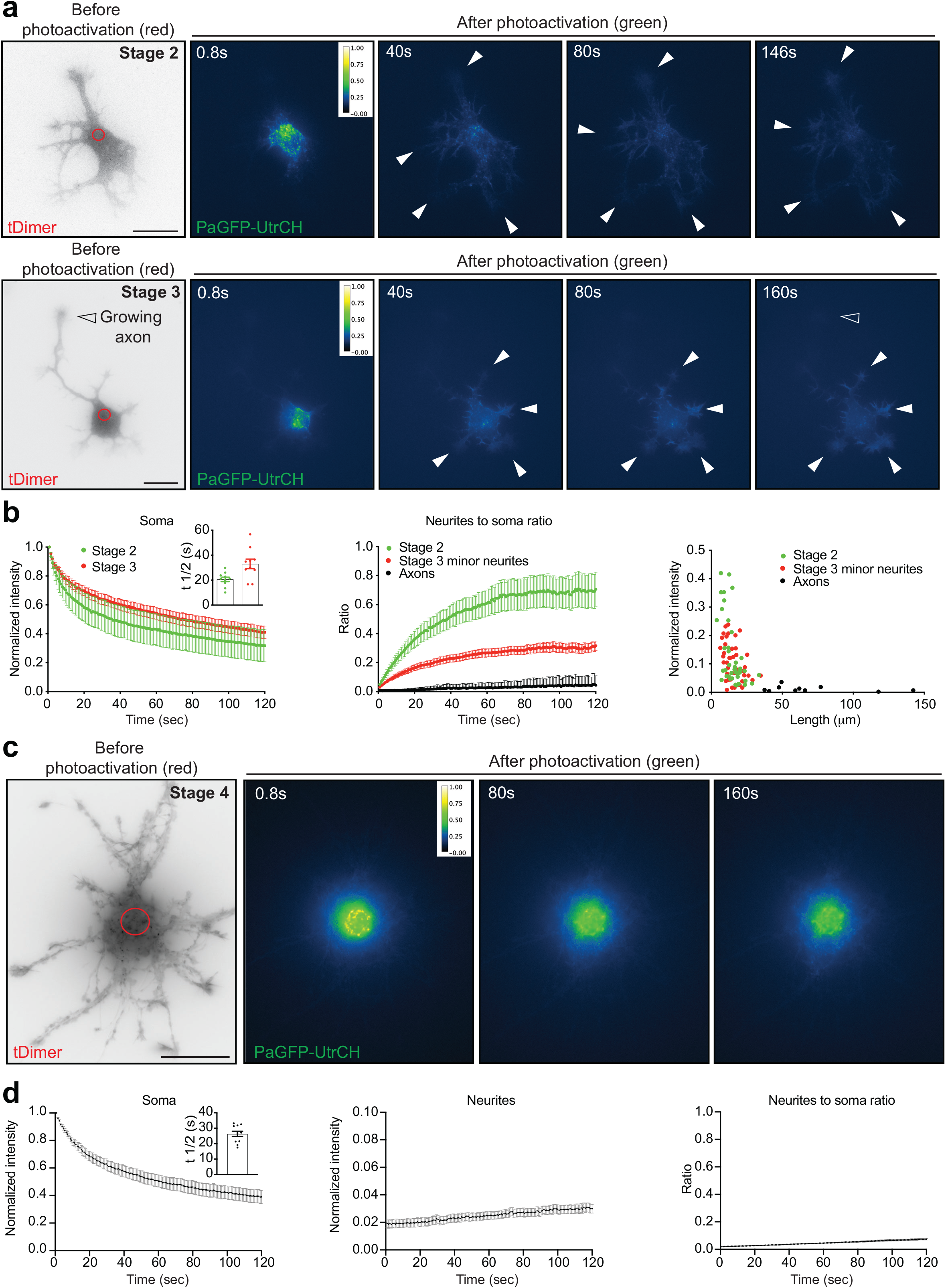
Radial somatic F-actin is specifically delivered to shorter neurites in developing stage 2 and stage 3 neurons but not to longer neurites or neurites in matured DIV8 neurons. a. PaGFP-UtrCH and tDimer co-transfected stage 2 (upper row) and stage 3 (lower row) rat hippocampal neurons photoactivated in the soma with a 405 nm laser (red circle with a diameter of 5.239 μm). b. Left panel: normalized intensity values in the photoactivated area of PaGFP-UtrCH expressing stage 2 and stage 3 cells. Inset graph: mean ± SEM half-time (t½) values in seconds: stage 2 = 20.60 ± 1.819; stage 3 = 33.02 ± 3.862. Middle panel: photoactivated signal in neurite tips over time relative to the average initial signal from the illuminated area of PaGFP-UtrCH expressing stage 2, stage 3 cells. Right panel: quantification shows photoactivated somatic PaGFP-UtrCH/F-actin normalized intensity enriched in the neurite tips against the length of their corresponding neurites. n = 10 per each stage, cells were obtained from at least three different cultures. c. PaGFP-UtrCH and tDimer co-transfected DIV8 (stage 4) rat hippocampal neuron photoactivated in the soma with a 405 nm laser (red circle with a diameter of 5.239 μm). d. Left panel: normalized intensity values in the photoactivated area of PaGFP-UtrCH expressing DIV8 (stage 4) cells. Inset graph: mean ± SEM half-time (t½) values in seconds = 26.28 ± 1.711. Middle panel: photoactivated signal in neurite shafts at 20 microns away from the illuminated area center over time relative to the average initial signal from the illuminated area of PaGFP-UtrCH expressing stage 4 cells. Right panel: Neurite shaft to soma photoactivated signal intensity ratio of PaGFP-UtrCH. n = 11 cells from at least three different cultures.

**Figure 2.**
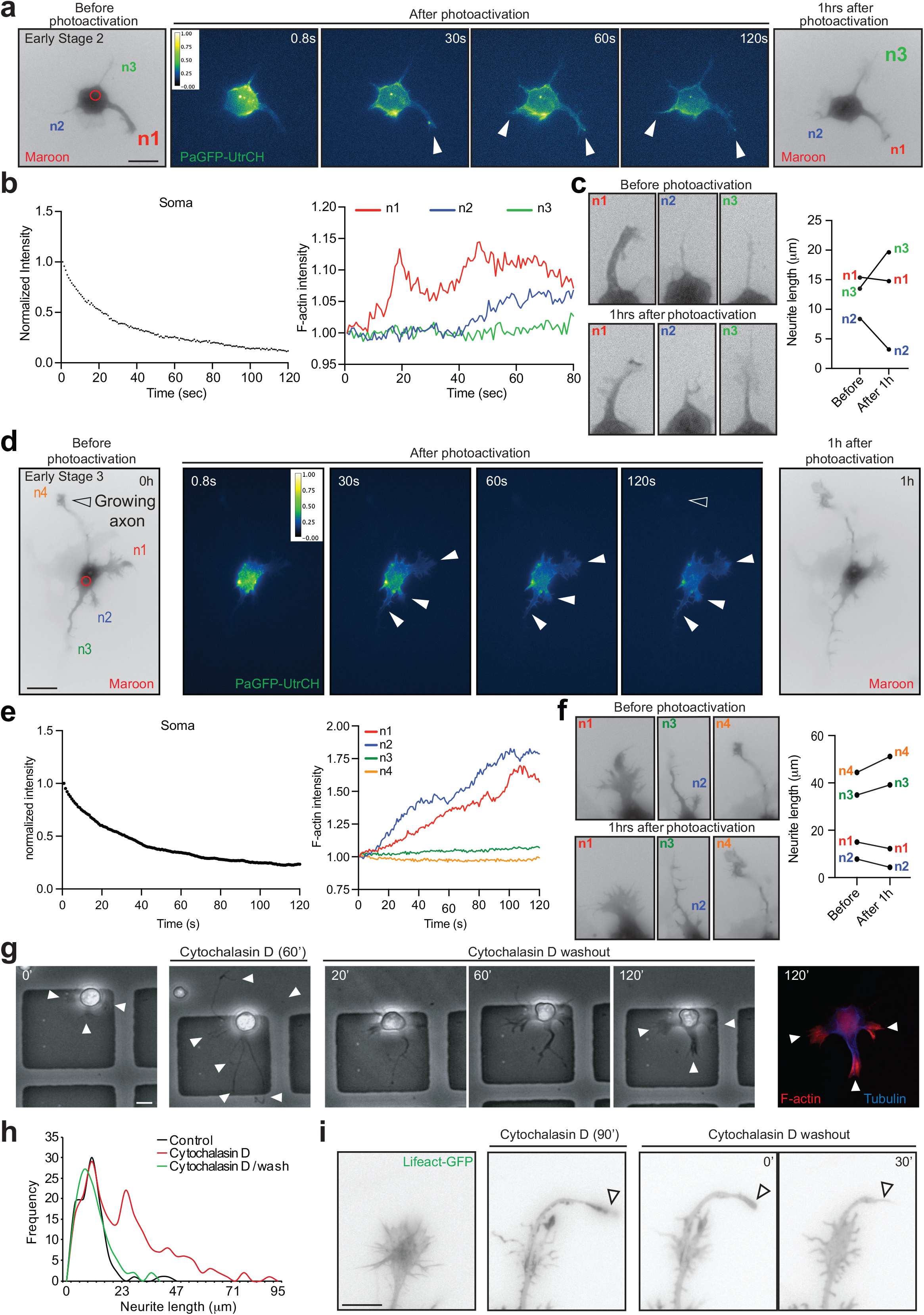
Somatic F-actin delivery negatively regulates neurite growth, and accordingly disruption of F-actin promotes neurite growth. a. PaGFP-UtrCH and mMaroon1 co-transfected stage 2 rat hippocampal neuron photoactivated in the soma with a 405 nm laser (red circle with a diameter of 5.239 μm), mMaroon1 channel imaged again 1h after photoactivation to track neurite morphology changes. b. Left panel: normalized intensity values in the photoactivated area of PaGFP-UtrCH expressing stage 2 cells. Right panel: photoactivated signal in neurite tips from 3 neurites, n1, n2, and n3 over time relative to the average initial signal from the neurite tips. c. The changes in neurite length before and 1h after photoactivation are shown in the cropped images of the neurites of the cell shown in a and the paired analysis graph. d. PaGFP-UtrCH and mMaroon1 co-transfected early stage 3 rat hippocampal neuron photoactivated in the soma with 405 nm laser (red circle with a diameter of 5.239 μm), mMaroon1 channel imaged again 1h after photoactivation to track neurite morphology changes. e. Left panel: normalized intensity values in the photoactivated area of PaGFP-UtrCH expressing early stage 3 cells. Right panel: photoactivated signal in neurite tips from 4 neurites, n1, n2, n3 and n4 over time relative to the initial signal. f. The changes in respective neurite length before and 1h after photoactivation are shown in the cropped images of the neurites of the cell shown in d and the paired analysis graph. g. Stage 2 cell treated for 60min with 2µM Cytochalasin D undergoes neurite elongation, which is normalized after 120min of washout, post hoc immunostained with F-actin (phalloidin-568) and Tubulin (blue). h. Quantification shows neurite outgrowth in Cytochalasin D condition compared to Control and Cytochalasin D/washout conditions. I. Images show neurite growth cone of Lifeact-GFP transfected DIV 1 hippocampal neuron through the course of 90min Cytochalasin D treatment and followed by washout. Kymographs with respective timestamps depict the loss of retrograde F-actin flow upon Cytochalasin D treatment and recovery followed by washout.

To further test this idea, we decided to affect F-actin dynamics using cytochalasin D (CytoD), which is known to induce neurite growth due to F-actin dynamics disruption at growth cones ^11^. We performed a time-lapse analysis to monitor neurite growth on stage 2 neurons expressing Lifeact-GFP to track F-actin behavior during and after CytoD treatment. As expected, CytoD treatment induced neurite growth (Figure 2g-i). Once CytoD was removed, however, F-actin reorganized in the neurite tips provoking neurite retraction (Figure 2g-i). Altogether, these results support the idea that somatic F-actin translocation into growth cones is an inhibitory signal for neurite growth.

### Myosin II mediates somatic F-actin translocation into neurite growth cones

To understand mechanistically how the somatic F-actin translocates into growth cones, we analyzed whether Myosin II is the molecular motor transporting radially the somatic F-actin. It is shown that during cytokinesis localization of F-actin is Myosin II-dependent ^25^. Moreover, in migrating cells Myosin II inhibition blocks actin network flow ^26^. Therefore, Myosin II is a suitable candidate to test whether this motor protein mediates somatic F-actin translocation into neurites. To investigate the role of Myosin II on somatic F-actin transport, first, we analyzed whether somatic PaGFP-Myosin IIA moves radially into neurites after photoactivation. We examined stage 2-early stage 3 neurons and we found that photoactivated PaGFP-Myosin IIA translocated from the soma into neurites in a similar fashion as photoactivated PaGFP-UtrCH with a preference to move into shorter neurites (Figure 3a-c; Video 4). Moreover, inhibition of Myosin II with blebbistatin reduced the photoactivated somatic PaGFP-UtrCH translocation into neurites (Figure 3d-g and Supp Fig 1; Video 5). Importantly, although blebbistatin affected somatic F-actin translocation into neurites, still the reduced photoactivated somatic PaGFP-UtrCH signal that moved into neurites following an inverse correlation with neurite length (Figure 3e). These results suggest that Myosin II mediates somatic F-actin translocation into neurites but the preference for somatic F-actin transport into minor neurites is not inherent to the Myosin II motor.

**Figure 3.**
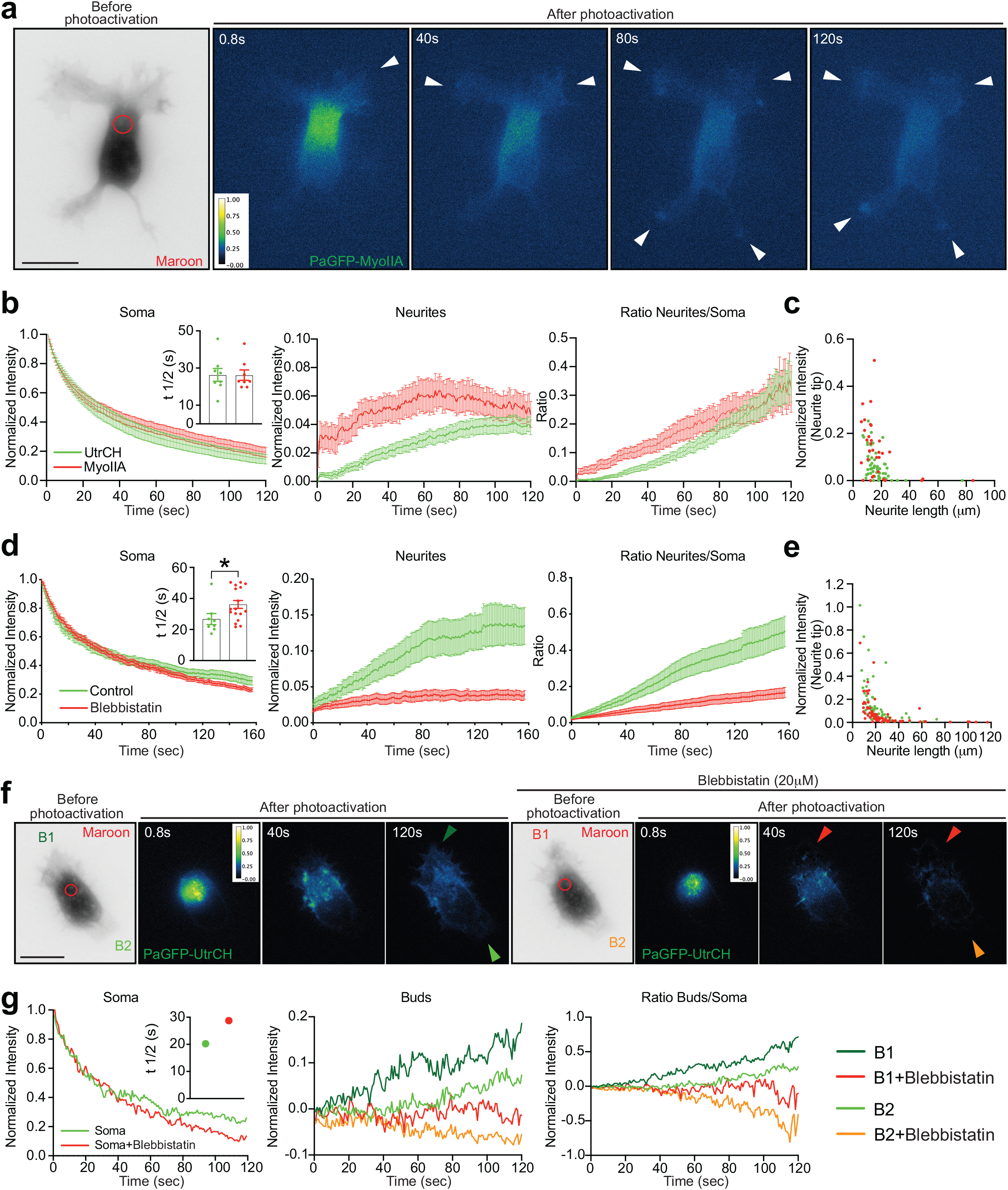
Similar to F-actin, somatic Myosin IIA follows radial delivery, and F-actin radial delivery is Myosin II-dependent. a. PaGFP-Myosin IIA and mMaroon1 co-transfected stage 2 rat hippocampal neuron photoactivated in the soma with a 405 nm laser (red circle with a diameter of 5.239 μm). b. Left panel: normalized intensity values in the photoactivated area of PaGFP-UtrCH and PaGFP-Myosin IIA expressing DIV1(stage 2) cells. Inset graph: mean ± SEM half-time (t½) values in seconds PaGFP-UtrCH = 26.32 ± 3.447 and PaGFP-Myosin IIA = 26.19 ± 2.801. Middle panel: photoactivated signal in neurite tips relative to the average initial signal from the illuminated area of PaGFP-UtrCH/Myosin IIA expressing DIV1(stage 2) cells. Right panel: neurite shaft to soma photoactivated signal intensity ratio of PaGFP-UtrCH/Myosin IIA. c. Quantification shows photoactivated somatic F-actin and Myosin IIA normalized intensity enriched in the neurite tips against the length of their corresponding neurites. For data shown in b and c, n = 8 cells per each group from at least three different cultures. d. Left panel: normalized intensity values in the photoactivated area of untreated Control and 20µM Blebbistatin-treated DIV1 (cells in Supp Figure 1) PaGFP-UtrCH expressing cells. Inset graph: mean ± SEM half-time (t½) values in seconds: Controls = 26.73 ± 3.534 and Blebbistatin cells = 36.11 ± 2.548. Middle panel: photoactivated signal in neurite tips relative to the average initial signal from the illuminated area for Control and 20µM Blebbistatin-treated DIV1 PaGFP-UtrCH expressing cells. Right panel: neurite shaft to soma photoactivated signal intensity ratio of PaGFP-UtrCH for Control and 20µM Blebbistatin-treated DIV1 PaGFP-UtrCH expressing cells. e. Quantification shows photoactivated somatic F-actin in Control and Blebbistatin-treated cells normalized intensity enriched in the neurite tips against the length of the neurites from which they are obtained. For data shown in d and e, n = 17 Control and 18 Blebbistatin-treated cells from at least two different cultures. f. PaGFP-UtrCH and mMaroon1 co-transfected stage 1 rat hippocampal neurons photoactivated in the soma with a 405 nm laser (red circle with a diameter of 5.239 μm) before and 30min after 20µM Blebbistatin treatment. B1 and B2 indicated on the stage 1 cell are the buds quantified. g. Left panel: normalized intensity values in the photoactivated area before and 30 min after 20µM Blebbistatin treatment. Inset graph: half-time (t½) values in seconds: Before = 20.22 and Blebbistatin treatment= 28.74. Middle panel: photoactivated signal in buds relative to the average initial signal from the illuminated area before and 30 min after 20µM Blebbistatin treatment. Right panel: buds to soma photoactivated signal intensity ratio before and 30 min after 20µM Blebbistatin treatment.

### Phosphorylation of actin at Tyr-53 in neurite shafts precludes the transport of somatic F-actin

To elucidate the mechanism by which there is a preferential somatic F-actin translocation into minor neurites but not the axon on stage 3 neurons, we decided to test whether the F-actin cytoskeleton instability plays a role. Given that Myosin II is an actin-directed motor, it is possible to envision that Myosin II moves along those actin filaments to transport somatic F-actin into neurites. Moreover, unstable actin filaments in the neurite shaft might preclude proper motility of somatic Myosin II into growth cones. In this regard, phosphorylation of actin at Tyr-53 (pY53) destabilizes actin filaments or F-actin ^17^. Therefore, we decided to characterize the distribution of actin pY53 in relation to somatic F-actin delivery in early developing neurons. To this end, we prepared hippocampal pyramidal cultured neurons at stages 2-3 for immunostaining to label pY53 actin. Importantly, we found that pY53 actin is enriched in the longest neurite shaft from stage 2 neurons, in the growing axon of stage 3 neurons (Figure 4a-d), and dendrites and axon of stage 4 neurons (Supp. Fig. 2). Consequently, photoactivated somatic PaGFP-UtrCH translocation into neurites was precluded from the neurite shaft with enriched pY53 actin signal (Figure 4e-h; Video 6) or from growing neurites (Figure 1). Interestingly, blebbistatin treatment, which induced increased neurite length, also increased the pY53 actin signal in those long-blebbistatin-induced neurites (Supp Fig 3). These data suggest that neurite growth correlates with pY53 actin.

**Figure 4.**
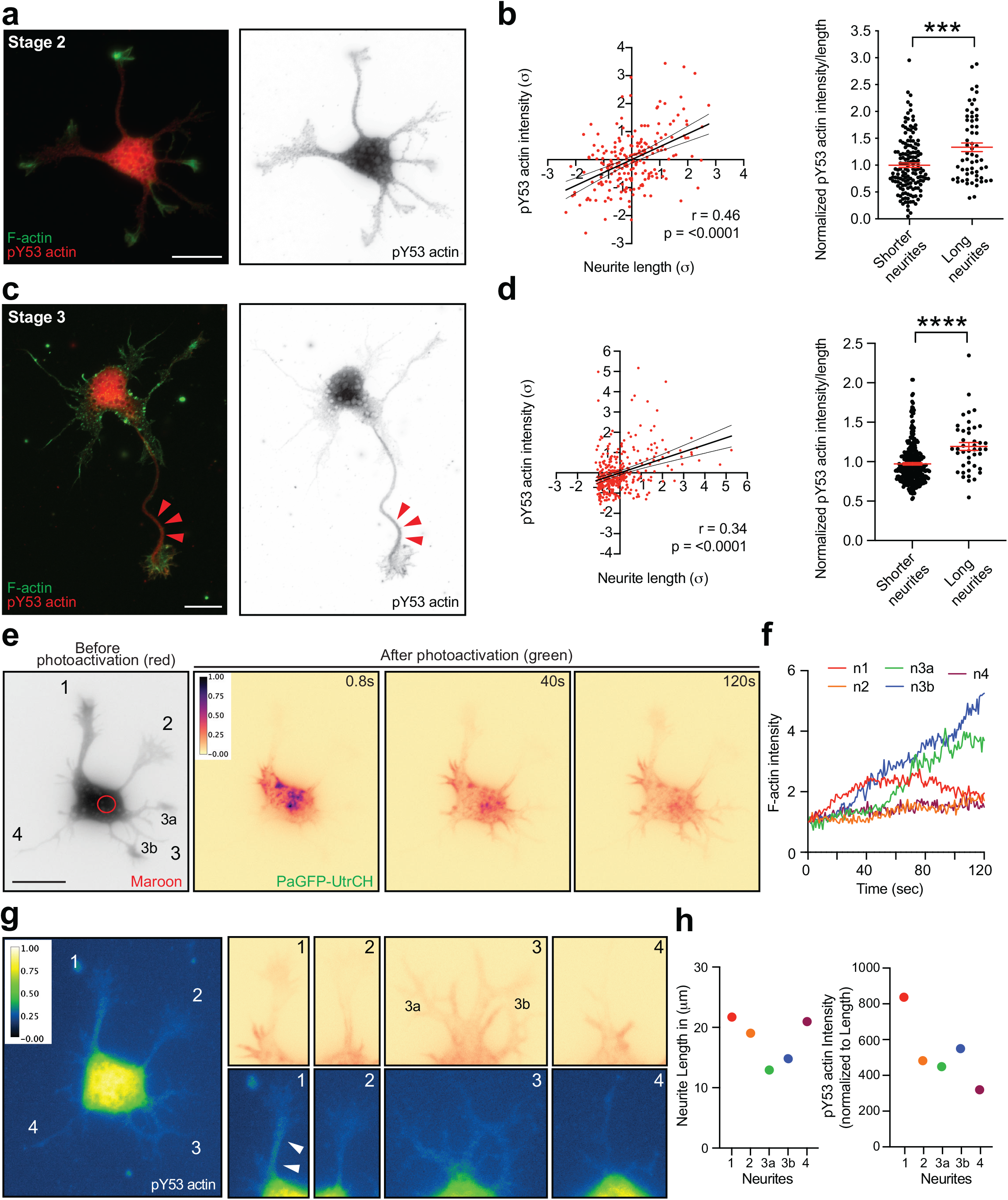
pY53 actin is enriched in the shaft of the longest neurite in developing neurons. a. Stage 2 neuron immunostained with pY53 actin and F-actin (labelled by phalloidin-488) b. Left panel: linear regression analysis of neurite length and pY53 actin intensity (normalized to Length) in stage 2 cells; values were normalized according to a standard score, and axes are represented in units of SD (σ). Linear regression line equation: Y = 0.4612*X - 9.973e-006. Pearson correlation coefficient (r value) and p value of significance are as indicated. Thin lines denote 95% confidence intervals. Right panel: quantification of pY53 actin intensity values normalized to length from shorter neurites and longer neurites (per each cell) at stage 2. Mean ± SEM values for normalized pY53 intensities in shorter neurites = 1.000 ± 0.04169 and long neurites = 1.333 ± 0.08102. *** P = 0.0006 by Mann-Whitney non-parametric test. Data were obtained from 210 neurites of 51 neurons from three different cultures. c. Stage 3 neuron immunostained with pY53 actin and F-actin (labelled by phalloidin-488) d. Left panel: linear regression analysis of neurite length and pY53 actin intensity (normalized to Length) in stage 3 cells; values were normalized according to a standard score, and axes are represented in units of SD (σ). Linear regression line equation: Y = 0.3452*X - 2.568e-007. Pearson correlation coefficient (r value) and p value of significance are as indicated. Thin lines denote 95% confidence intervals. Right panel: quantification of pY53 actin intensity values normalized to length from shorter neurites and longer neurites (per each cell) at stage 3. Mean ± SEM values for normalized pY53 intensities in shorter neurites = 0.9725 ± 0.01504 and longer neurites = 1.192 ± 0.05046. **** P <0.0001 by Mann-Whitney non-parametric test. Data were obtained from 344 neurites of 43 neurons from three different cultures. e. PaGFP-UtrCH and mMaroon1 co-transfected stage 2 rat hippocampal neuron photoactivated in the soma with a 405 nm laser (red circle with a diameter of 5.239 μm) f. Photoactivated signal enriched in the neurite tips from 4 neurites n1, n2, and n3a, n3b and n4 over time relative to the initial signal (shown in e). g. Stage 2 neuron from e, post hoc immunostained with pY53 actin. Cropped images from all 4 neurites show an enrichment of photoactivated signal (upper panels) and pY53 immunostaining (lower panels). h. Graphs depicting the length of individual neurites before photoactivation from mMaroon1 channel shown in e (left) and pY53 actin intensity (normalized to Length) values of individual neurites after photoactivation followed by post hoc pY53 actin immunostaining.

To directly test the relevance of actin phosphorylation at Y53 in neurite outgrowth/neuronal polarization, we decided to introduce the phospho-mimetic (Y53E) and phospho-dead (Y53A) forms of pY53 actin ^16^. To do so, we transfected neurons before plating with wt actin-mCherry, Y53E actin-GFP, and Y53A actin-mCherry. Two days after plating neurons were fixed and prepared for immunostaining to label growing axons with the axonal marker Tau-1. Our results showed that the preclusion of actin phosphorylation (Y53A) reduced the number of neurons extending an axon compared with neurons expressing wt actin (Supp. Fig. 4a, c). On the contrary, constitutive phosphorylation of actin (Y53E) induced the formation of longer axons, compared with neurons expressing wt actin (Supp. Fig. 4a, b). Altogether, these results support the idea that pY53 actin promotes neurite growth.

To learn more about the molecular characteristics of the neurites enriched with pY53 actin, we used truncated Kinesis-1 (Kif5c), which is shown to accumulate transiently in neurites of stage 2 neurons and only in the emerging axon of stage 3 neurons ^27^. We transfected neurons before plating with Kif5c-Td tomato plasmid and 1-2 days after plating cells were fixed and prepared for immunostaining to label pY53 actin. Our results show that the neurite shaft with increased pY53 actin signal is the one bearing the Kif5c-Td tomato signal in stage 2 and 3 neurons (Figure 5). Moreover, we found that the neurites exhibiting Kif5c-Td tomato also have increased acetylated tubulin (Figure 5) as previously shown ^28^. Finally, we performed photoactivation of somatic PaGFP-UtrCH in stage 2 cells co-transfected with Kif5c-Td tomato. We found that photoactivated somatic PaGFP-UtrCH translocation occurred preferentially in neurites lacking the Kif5c-Td tomato signal (Figure 5e-g; Video 7). Thus, suggesting that somatic F-actin translocate into neurites with less pY53 actin.

**Figure 5.**
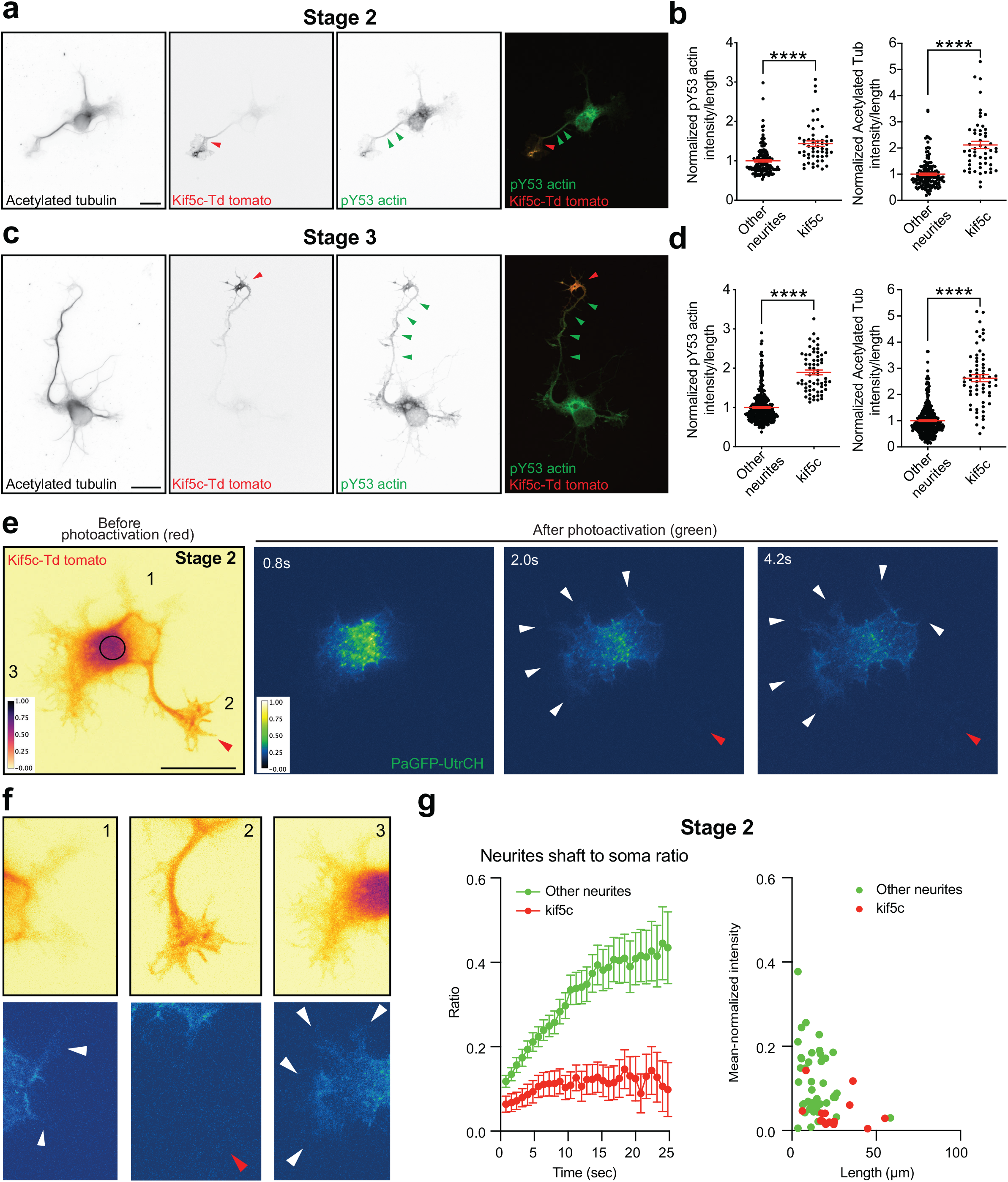
pY53 actin and Acetylated tubulin are enriched in the shaft of neurites bearing Kif5c. a. Images for Kif5c-Td tomato transfected stage 2 neurons immunostained with pY53 actin and AcetyTUB. b. Left panel: quantification of pY53 actin intensity values normalized to length from neurites and Kif5c-positive neurites (per each cell) at stage 2. Mean ± SEM values for normalized pY53 intensities in shorter neurites = 1.000 ± 0.02989 and long neurites = 1.437 ± 0.07009. **** P <0.0001 by Mann-Whitney non-parametric test. Right panel: quantification of AcetyTUB intensity values normalized to length from neurites and Kif5c-positive neurites (per each cell) at stage 2. Mean ± SEM values for normalized AcetyTUB intensity in shorter neurites = 1.000 ± 0.04479 and long neurites = 2.119 ± 0.1409. **** P <0.0001 by Mann-Whitney non-parametric test. Data were obtained from 198 neurites of 46 neurons from three different cultures. c. Images for Kif5c-Td tomato transfected stage 3 neurons immunostained with pY53 actin and AcetyTUB. d. Left panel: quantification of pY53 actin intensity values normalized to length from neurites and Kif5c-positive neurites (per each cell) at stage 3. Mean ± SEM values for normalized pY53 intensities in shorter neurites = 1.000 ± 0.02135 and long neurites = 1.893 ± 0.06266. **** P <0.0001 by Mann-Whitney non-parametric test. Right panel: quantification of AcetyTUB intensity values normalized to length from neurites and Kif5c-positive neurites (per each cell) at stage 3. Mean ± SEM values for normalized AcetyTUB intensity in shorter neurites = 1.000 ± 0.03061 and long neurites = 2.624 ± 0.1325. **** P <0.0001 by Mann-Whitney non-parametric test. Data were obtained from 415 neurites of 59 neurons from three different cultures. e. PaGFP-UtrCH and Kif5c-Td tomato co-transfected stage 2 rat hippocampal neuron photoactivated in the soma with a 405 nm laser (red circle with a diameter of 5.215 μm). f. Cropped images of the neurites of the cell shown in e. g. Left panel: photoactivated signal in neurite shafts at 12 microns away from the illuminated area center over time relative to the average initial signal from the illuminated area of PaGFP-UtrCH expressing stage 2 cells. Right panel: quantification shows photoactivated somatic F-actin mean-normalized intensity enriched in the neurite tips against the length of their corresponding neurites. n = 15 cells per each group from four different cultures.

### Acetylated microtubules favor the enrichment of actin phosphorylation at Tyr-53 in the neurite shaft

Next, we decided to investigate determinants of pY53 actin enrichment in the neurite shaft. It has been reported an association of actin filaments with microtubules in neurites shaft ^29^. Moreover, that acetylated tubulin is enriched in the shaft of the growing axon ^15, 18^. Therefore, we asked whether the localization of acetylated tubulin and pY53 actin coexist in the same neurite shaft domain. Using immunofluorescence, we found that there was a positive correlation between acetylated tubulin and pY53 actin signals in the neurite shaft of stage 2 and 3 neurons, specifically in the growing axon of stage 3 neurons (Figure 6a-d). We obtained similar results for polyglutamylated tubulin (Supp. Fig. 5). Furthermore, taxol treatment, which increases neurite length and tubulin acetylation ^15, 18^ in the neurite shaft, induced the increased pY53 actin signal in neurites (Figure 6e-g). On the other hand, though polyglutamylated tubulin levels increased upon Taxol treatment, its disribution showed a weaker coorelation with that of pY53 actin (Sup Figure 5 c, d) compared to the acetylated tubulin (Figure 6g, i). Importantly, however, a short period of treatment with taxol, which did not affect neurite length still increased tubulin acetylation in the neurites together with increased pY53 actin signal in those neurite shafts (Figure 6h, j). This data suggest that the pY53 actin is not completely length-dependent but more directly influenced by tubulin acetylation. Nocodazole treatment, which induces microtubule depolymerization, reduced the levels of pY53 actin (Supp. Fig. 6). Collectively, our results show that pY53 actin precludes somatic F-actin translocation into neurites and that this actin modification is influenced by the acetylation of tubulin, ultimately leading to timely neurite growth, first the axon and then the dendrites.

**Figure 6.**
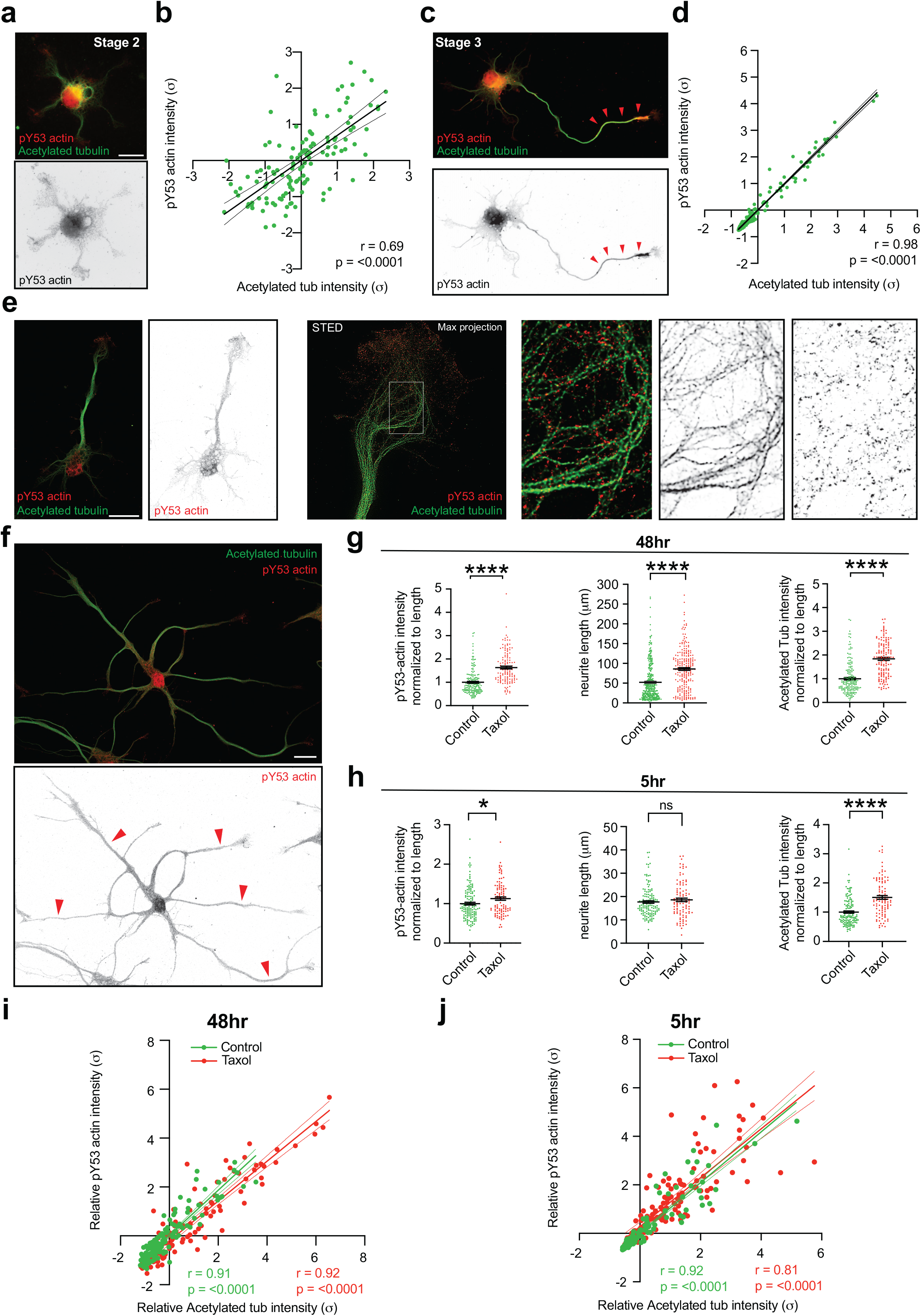
pY53 actin distribution and acetylated tubulin profile correlate strongly in the neurite shafts of developing neurons. a. Stage 2 neuron immunostained with pY53 actin and AcetyTUB. b. Linear regression analysis of AcetyTUB and pY53 actin intensities in stage 2 cells; values were normalized according to a standard score, and axes are represented in units of SD (σ). Linear regression line equation: Y = 0.6963*X + 7.634e-008. Pearson correlation coefficient (r value) and p value of significance are as indicated. Thin lines denote 95% confidence intervals. Data were obtained from 128 neurites of 49 neurons from two different cultures. c. Stage 3 neuron immunostained with pY53 actin and AcetyTUB. d. Linear regression analysis of AcetyTUB and pY53 actin intensities in stage 3 cells; values were normalized according to a standard score, and axes are represented in units of SD (σ). Linear regression line equation: Y = 0.9852*X - 2.720e-007. Pearson correlation coefficient (r value) and p value of significance are as indicated. Thin lines denote 95% confidence intervals. Data were obtained from 175 neurites of 30 neurons from three different cultures. e. STED images of the growth cone of a stage 3 neuron immunostained with pY53 actin and AcetyTUB, inset from the white rectangle shows AcetyTUB and pY53 actin signals overlap. f. Rat hippocampal neuron treated with 5nM Taxol (24h after plating) for 48h, immunostained with pY53 actin and AcetyTUB. g. Quantifications from Control and Taxol-treated (48h). Left and Right panels: Data were obtained from 175 neurites from 30 Control cells and 134 neurites from 30 Taxol-treated cells from three different cultures. Middle panel: Data were obtained from 344 neurites from 43 Control cells and 229 neurites from 43 Taxol-treated (48h) cells from three different cultures. Left panel: quantification of relative pY53 actin intensity values normalized to length from the neurite shafts. Mean ± SEM values from Control neurons = 1.000 ± 0.04077 and Taxol-treated (48h) neurons = 1.633 ± 0.05916. **** P <0.0001 by Mann-Whitney non-parametric test. Middle panel: quantification of neurite length. Mean ± SEM values from Control neurons = 52.05 ± 2.488 and Taxol-treated (48h) neurons = 85.76 ± 3.452. **** P <0.0001 by Mann-Whitney non-parametric test. Right panel: quantification of relative AcetyTUB intensity values normalized to length from the neurite shafts. Mean ± SEM values from Control neurons = 1.000 ± 0.05084 and Taxol-treated (48h) neurons = 1.839 ± 0.06441. **** P <0.0001 by Mann-Whitney non-parametric test. h. Quantifications from Control neurons and neurons incubated with 5nM Taxol (immediately after plating) for 5h. Data were obtained from 143 neurites from 64 control cells and 99 neurites from 52 Taxol-treated cells from two different cultures. Left panel: quantification of pY53 actin intensity values normalized to length from the neurite shafts. Mean ± SEM values from control neurons = 1.000 ± 0.03340 and Taxol-treated (5h) neurons = 1.136 ± 0.04187. ** P = 0.0100 by Mann-Whitney non-parametric test. Middle panel: quantification of neurite length. Mean ± SEM values from Control neurons = 17.67 ± 0.5277 and Taxol-treated (5h) neurons = 18.60 ± 0.7689. P = 0.3866 by Mann-Whitney non-parametric test, n.s: not significant. Right panel: quantification of AcetyTUB intensity values normalized to length from the neurite shafts. Mean ± SEM values from Control neurons = 1.000 ± 0.03860 and Taxol-treated (5h) neurons = 1.505 ± 0.06407. **** P <0.0001 by Mann-Whitney non-parametric test. i. Linear regression analysis of AcetyTUB and pY53 actin intensities in Control and Taxol groups (neurons incubated with 5nM Taxol 24h after plating for 48h); values were normalized according to a standard score, and axes are represented in units of SD (σ). Linear regression line equations for pY53 actin intensities from Control group: Y = 1.046*X + 1.704e-007; from Taxol group: Y = 1.008*X + 0.2777. Pearson correlation coefficient (r value) and p value of significance are as indicated. Thin lines denote 95% confidence intervals. Data were obtained from 175 neurites of 30 Control neurons and 134 neurites of 30 Taxol-treated neurons from three different cultures. j. Linear regression analysis of AcetyTUB and pY53 actin intensities in Control and Taxol-treated cells (neurons incubated with 5nM Taxol immediately after plating for 5h); values were normalized according to standard score, and axes are represented in units of SD (σ). Linear regression line equations for pY53 actin intensities from Control group: Y = 0.9235*X + 0.01825, from Taxol group: Y = 0.8150*X - 0.2079. Pearson correlation coefficient (r value) and p value of significance are as indicated. Thin lines denote 95% confidence intervals. Data were obtained from 143 neurites from 64 control cells and 99 neurites from 52 Taxol-treated (5h) cells from two different cultures.

## Discussion

During neuronal polarization of pyramidal neurons, one neurite grows as an axon meanwhile the others remain short. Eventually, once the axon is established, the other neurites initiate the outgrowth as dendrites. This pattern of differentiation is not only reported in cultured neurons^1^ but also *in vivo*/*in situ* ^2, 3, 20^. One important question in neurobiology is how these sequential steps of differentiation are achieved. It was suggested an intrinsic feedback system with inhibitory internal signals precluding neurite extension of future dendrites during axon elongation. On the other hand, positive molecular cascades might sustain locally the axon extension ^4^. Now, we show that somatic F-actin functions as a neurite growth inhibitor at the growth cones (Figure 2); specifically, at growth cones of the short neurites that do not grow during axon extension. Moreover, inhibition of Myosin II, hence disrupting somatic F-actin transport into growth cones, induces neurite extension of shorter neurites but not the ones growing as an axon (Supp Fig. 3c). To our knowledge, this is the first time that it is shown that neuronal differentiation is mediated by somatic F-actin translocation into growth cones. Specifically, we show that this specific transport restricts neurite extension and helps to support the formation of one axon and several dendrites in a sequential fashion. Thus, our results suggest that the internal inhibitory signal that precludes neurite extension during neuronal polarization is the somatic F-actin translocation into growth cones.

Mechanistically, we found that phosphorylation of actin at Tyr-53 (pY53) in the neurite shaft precludes somatic F-actin transport into the growing neurite (Figure 4). It is shown that this actin phosphorylation destabilizes actin filaments or F-actin ^17^. Therefore, unstable actin filaments in the neurite shaft might preclude proper motility of somatic Myosin II/F-actin into growth cones. Interestingly, the phospho-mimetic (Y53E) form of pY53 actin only induces neurons with longer axons and not neurons with multiple axons (Supp Fig. 4). This could be explained by the fact that initial axon formation is microtubule stability-dependent rather than actin cytoskeleton-mediated ^15^. It is shown that early-developing neurons are not capable of forming multiple axons if their actin cytoskeleton is disrupted, only if the microtubule stabilization is promoted ^15^. Accordingly, now we show that the pattern of actin phosphorylation (Y53) in the neurite shaft intermingles with acetylated microtubules (Figure 6). Thus, pharmacological induction of microtubule acetylation increases the pY53 actin signal in all neurites. Finally, during dendrites elongation, we found that somatic F-actin transport towards growing neurites is precluded. Hence, the pY53 actin signal is increased in growing dendrites (Supp Fig. 2). Overall, our data suggest that neurite outgrowth is supported by actin phosphorylation (Y53) in the neurite shaft, which in turn precludes somatic F-actin translocation into the growth cones.

Finally, we found that truncated Kinesin-1 (Kif5c), which is shown to accumulate transiently in neurites of stage 2 neurons and only in the emerging axon of stage 3 neurons ^27^, localizes at the growth cone of the neurite with increased pY53 actin and acetylated tubulin signal (Figure 5). In this regard, it is shown that microtubule acetylation promotes Kinesin-1 binding and transport ^28^. Given that in stage 2 neurons the neurites grow and retract until one keeps growing as an axon (stage 3)^1^, the actin phosphorylation (Y53) in the neurite shaft may be a dynamic process until one is selected to grow as an axon. Nevertheless, our data do not explain how actin phosphorylation takes place in growing neurites. Our results, however, allow us to envision that microtubule stability might be the triggering factor mediating this specific actin phosphorylation in the growing neurite shaft. More experiments are required to explore this possibility.

## Material and methods

### Plasmid constructs

Thomas Oertner (ZMNH, UKE) kindly provided the tDimer (pAAV-CAG-tDimer) plasmid. mMaroon1 (pcDNA3.1-mMaroon1) plasmid was provided by Michael Lin (Addgene plasmid # 83840; http://n2t.net/addgene: 83840; RRID: Addgene_83840; ^30^). mRuby2-Actin-C-18 was a gift from Michael Davidson (Addgene plasmid # 55889; http://n2t.net/addgene:55889; RRID: Addgene_55889). PaGFP-UtrCH was a gift from William Bement (Addgene plasmid # 26738; http://n2t.net/addgene:26738; RRID: Addgene_26738; ^24^). mPA-GFP-MyosinIIA-C-18 was a gift from Michael Davidson (Addgene plasmid # 57149; http://n2t.net/addgene:57149; RRID: Addgene_57149). pBa-KIF5C 559-tdTomato-FKBP was a gift from Gary Banker (Addgene plasmid # 64211; http://n2t.net/addgene:64211; RRID: Addgene_64211). mCherry-β-actin, mCherry-Y53A-β-actin and GFP-Y53E-β-actin constructs were kindly provided by Pirta Hotulainen ^16^.

### Rat primary hippocampal neuron cultures and transfections

Pregnant rats were anesthetized with CO2/O2 and euthanized before taking the E18 embryos out from their uteri. Embryos were then decapitated, skulls were opened, and brains were collected in petri dishes with HBSS on ice. Hemispheres were separated, meninges were carefully stripped away, and hippocampi were dissected on ice and triturated in 1xHBSS (Invitrogen) after digestion by papain and DNase (Worthington for 10 min at 37°C). Transfections were performed using the Amaxa nucleofector system following the manufacturer’s manual. The final concentration for the Actin-mRuby, mMarron1, PaGFP-UtrCH and PaGFP-MyosinIIA was 1 µg. Empty pcDNA 3.1 was used to make up to 3 µg of DNA for 5 × 10^6^ cells per each transfection mix as per the manufacturer’s recommendation. After electroporation, neurons were plated on poly-L-lysine coated coverslips or tissue culture chambers (Sarstedt, for live imaging) in Neurobasal/B27 medium (Invitrogen) and were maintained in culture for 24 to 72h or 7-8 days at 37°C with 5% CO2 before use.

### Photoactivation experiments

Photoactivation experiments of PaGFP-UtrCH and PaGFP-MyosinIIA were performed in the soma of primary rat hippocampal neurons using either the iLAS2 – Visitron setup (Figures 1, 3d-g, Supp Figures 1, described earlier ^10^) or NIKON FRAP setup.

On the iLAS2 – Visitron setup the time-lapses were imaged with a 100x TIRF objective (Nikon, ApoTIRF 100X oil (NA 1.49). Emission light was collected through a quad-band filter (Chroma, 405/488/561/640) followed by a filter wheel with filters for GFP (Chroma, 525/50 m), RFP (Chroma, 595/50 m), and Cy5 (Chroma, 700/75 m). Multichannel images were acquired sequentially with an Orca flash 4.0LT CMOS camera (Hamamatsu) controlled by VisiView software (Visitron Systems Gmbh).

On the NIKON FRAP system, images were taken with PLAN APO 60X oil (NA 1.42) objectives on a Nikon EclipseTi2 inverted microscope equipped with a SPECTRA X Light Engine as an LED light source (Lumencor^®^ from AHF analysentechnik AG, Germany), and a digital CMOS camera ORCA-Flash4.0 V3 C13440-20CU (Hamamatsu, Japan) controlled with NIS-Elements software.

During the time-lapse imaging, cells plated on 4 well culture chambers (Sarstedt or Ibidi) were kept in an acrylic chamber at 37°C in 5% CO2. To identify transfected neurons before photoactivation, the cultures used for this experiment were always co-transfected with Actin-mRuby or mMaroon1 together with PaGFP-tagged constructs. To illustrate the neuronal morphology, an image with Actin-mRuby or mMaroon1 signal was captured before the photoactivation with 561 nm or 640 nm lasers, respectively.

Photoactivation was achieved using the 405 nm laser with 50% power (iLAS2 system) or 10% power (NIKON FRAP system). The diameter of the 405 nm laser illuminated area for all the cells used for the analysis was either 5.239 µm on the iLAS2 system or 5.239 µm on the NIKON FRAP system; the 405 nm laser dwell time was 2 ms per pixel (1 pixel = 65 nm) on iLAS2 system or 600 µs per pixel (1 pixel = 73.4 nm) on NIKON FRAP system. Cells were illuminated with 488 nm 5% laser on the iLAS2 system or 10% LED power on the NIKON FRAP system for 800 ms to record one frame. Three frames were imaged before, and 150 or 200 frames were recorded after the photoactivation.

### Photoactivation analysis

In every time-lapse, photobleaching was checked and, if applicable, corrected with Exponential Bleach (Fiji-ImageJ, NIH). For analysis, the area of photoactivation at either soma was selected as an ROI. Further ROIs were selected in the corresponding compartments (soma, neurite tip). For the soma, we selected the region of photoactivation, for growth cones/neurite tips we selected an area, which remains within the central region, even during eventual movement. If the growth cone was moving too much, the region was cropped and aligned via ImageJ’s template matching plugin to avoid moving artefacts. Background/autofluorescence signal was calculated from the average grey values of the first three frames that were imaged before the photoactivation, these values were then subtracted from the grey values after photoactivation to plot the graphs. For plots, 0 was considered as the first frame after photoactivation. Average grey values over time were measured via the Time series Analyzer plugin. Initial grey values in the photoconverted area were normalized to 1; grey values in the “receiving” compartments (neurite tips) were normalized to fractions of the initial grey value at the photoconverted area (in the soma). The ratio of the average neurite tip intensity to the average soma intensity was plotted over time for different experimental groups. The decay of signal in the photoconverted area in the soma t½ was determined via Graph-Pad’s fitted one-phase exponential decay equation.

### *In utero* electroporation (IUE)

Pregnant C57BL/6 mice with E13 or E15 embryos were first administered with pre-operative analgesic, buprenorphine (0.1 mg/kg), by subcutaneous injection. After 30 min, mice were anesthetized with isoflurane (4% for induction, 2–3% for maintenance) in oxygen (0.5–0.8 l/ min for induction and maintenance). Later, uterine horns were exposed, and plasmids mixed with Fast Green (Sigma) were microinjected into the lateral ventricles of embryos. We introduced mCherry-β-actin, mCherry-Y53A-β-actin and GFP-Y53E-β-actin plasmids in combination with Venus or tDimer plasmids into brain cortices at E15. We used 1 µg/µl for the β-actin plasmids constructs and 0.5 µg/µl of tDimer or Venus. Five current pulses (50 ms pulse/950 ms interval) at 35V for E15 were delivered across the heads of embryos. After surgery, mice were kept in a warm environment and were provided with moist food containing post-operative analgesic, meloxicam (0.2-1 mg/kg), until they were euthanized for preparing primary cortical cultures.

### Primary mouse cortical cultures

Two days later pregnant mice were anesthetized with CO2/O2, and euthanized before taking the E17.5 embryos out from their uteri. Embryos were then decapitated, skulls were opened, and brains were collected in petri dishes with Hibernate-E medium (Invitrogen) on ice. Hemispheres were separated, meninges were carefully stripped away, and cortices were dissected on ice. Transfected (fluorescent) cortical regions were identified and dissected on ice under a stereo microscope (Olympus SZX16) equipped with a UV light source. The isolated cortical regions were first incubated in 1x HBSS (Invitrogen) with papain and DNase (Worthington) for 10 min at 37°C neurons and then triturated. The cells were then pelleted and washed with fresh HBSS before they were plated on poly-L-lysine coated coverslips or tissue culture chambers (Sarstedt, for live-imaging) in Neurobasal/B27 medium (Invitrogen), maintained in culture for 48 to 72h at 37°C with 5% CO2 before use for immunostainings.

### Pharmacological treatments

Cytochalasin D (Sigma, #C2618-200UL) - 2µM:

For data shown in Figure 2 g, h and i, DIV1 (14h to 18h old) neurons were treated for 60 or 90 min, washed out and fixed later.

S-nitro-Blebbistatin (Cayman chemicals, #856925-75-2) - 20µM:

For photoactivation experiments shown in Figures 3 d, f and g; Sup Figure 1b, DIV1 (14 to 18h old) neurons were treated for 15 to 75 min. For data shown in Sup Figure 3 a-f, neurons were treated at DIV1 (24h old) and fixed at DIV3.

Taxol (Sigma, #T7402) - 5nM:

For data shown in Figure 6 h and j, neurons were treated at the time of plating (at 0h) and fixed 5h later. For data shown in Figure 6 f, g, i and Supp figure 5c, d; neurons were treated at DIV1 (24h old) and fixed at DIV2 (48h after plating).

Nocodazole (Sigma, M1404-10MG) - 6µM:

For data shown in Supp Figure 6, DIV1 (24h old) neurons were treated for 30 min and fixed with 4% PFA for 2min and with 100% Methanol (pre-chilled at -20°C) for 3 minutes.

### Immunofluorescence

Rat hippocampal or mouse cortical neurons were grown on coverslips were fixed either with 4% paraformaldehyde (PFA) at 37°C for 10 min or with 4% paraformaldehyde (PFA) at 37°C for 2 min, followed by 3 min ice cold Methanol incubation at -20 °C. Cells were then permeabilized with 0.5% Triton X-100 for 10 min. Non-specific binding was blocked by incubation with 5% donkey serum in PBS for 60 min at RT (room temperature), followed by specific primary antibody incubation for 180 min at RT or overnight at 4°C, followed by 3 times 4 minutes PBS washes. Coverslips were incubated with respective anti-mouse or anti-rabbit Alexa Fluor 488 or 568 or 647 secondary antibodies, along with Phalloidin-488 (1:120, Cytoskeleton # PHDG1) to stain F-actin (only on PFA-fixed cells) and/or Hoechst dye (1:10,000, Invitrogen # 33258) to stain for nuclei for 60 min at RT followed by three washing steps with PBS. Coverslips were mounted onto slides using Fluoromount-G® (SouthernBiotech) and were stored and protected from light.

Antibodies used for immunostainings:

Primary – mouse anti-AcetyTub, 1:700 (Sigma #T7451); mouse anti-Tau-1, 1:700 (Millipore #MAB3420); Rat anti-α-Tubulin (SySy #302217), Rabbit anti-Phospho-Y53-actin (MyBioSource #MBS474080), Chicken anti-MAP2, 1:500 (SySy #188006). Mouse anti-Polyglutamylated tubulin (PolyGTUB) (Sigma #T9822).

Secondary – donkey anti-mouse IgG AF 488, 1:1000 (Invitrogen #A21202); donkey anti-mouse IgG AF 647, 1:1000 (Invitrogen #A31571); goat anti-rabbit IgG AF 488, 1:1000 (Invitrogen #A11077); donkey anti-rabbit IgG AF 647, 1:1000 (Invitrogen #A31573); anti-rat goat IgG AF 568, 1:1000 (Invitrogen #A11077).

### Fixed cell imaging

Images were taken on with a 60X oil (NA 1.42) objective on a Nikon EclipseTi2 inverted spinning disk confocal (X-Light V2 L-FOV from CrestOptics S.p.A. Italy) microscope equipped with a SPECTRA X Light Engine as an LED light source (Lumencor^®^ from AHF analysentechnik AG, Germany), and a digital CMOS camera ORCA-Flash4.0 V3 C13440-20CU (Hamamatsu, Japan) controlled with NIS-Elements software. LED power for 405, 488, 568 and 647 channels were set to 10%, with an exposure time ranging between 100-400 ms for all the fluorophores except for Hoechst dye (10 ms). Optical configuration settings were set the same between the experimental groups that were compared for analysis.

### Neurite analysis (intensity, length, and number)

pY53 actin and AcetyTUB intensity measurements in the neurite shaft: neurite shafts were traced with a segmented line option and saved to the ROI manager in Fiji from which the total intensity and length measurements were obtained. The ratio of the total intensities to length was performed to obtain normalized intensity values plotted in graphs.

Neurite length and neurite number analysis: neurites were traced with a segmented line option and saved to the ROI manager in Fiji from which the length measurements were obtained. For each cell, the length of all neurites, the length of the longest neurite and the length of other neurites were plotted. For neurite number analysis, the total number of neurites per cell was obtained from the ROIs obtained above.

### Image processing

Linear adjustment of brightness and contrast was performed on images using Photoshop CS or Fiji - ImageJ.

### Quantification and statistical analysis

Statistical analysis was performed using GraphPad Prism software versions 8 and 9. To compare the means of two groups, data were first tested for normal distribution and when the data passed the normality test, Student’s t-test (two-tailed) was used, otherwise its nonparametric counterpart Mann-Whitney test was used to compare the means of two groups. Asterisks *, **, *** and **** represent P < 0.05, 0.01, 0.001, and 0.0001, respectively. Error bars in the graphs always represent the standard error of the mean.

Pearson correlation analysis was performed for showing a linear relationship between two sets of data. Figure 4b (left panel), 4d (left panel); Figure 6b, 6d, and Supp Figures 5a, 5b the normalized intensity or neurite length values obtained from each neurite or shaft were first normalized to the mean value of all the shafts or neurites obtained from the corresponding cell. For data shown in Figures 6i, 6j, Supp Figure 3d and Supp Figure 5c the intensity values obtained from all neurite shafts in control and treatment groups were first normalized to the mean value of all the shafts measured in the control group in each experiment. For data shown in Supp Figure 2b, the normalized intensity values obtained from all neurite shafts were first normalized to the mean value of all the shafts measured. Finally, values were normalized according to the standard score (z-score) and axes are represented in units of standard deviation [σ].

## Supplementary Figure Legends

**Supp Figure 1.**
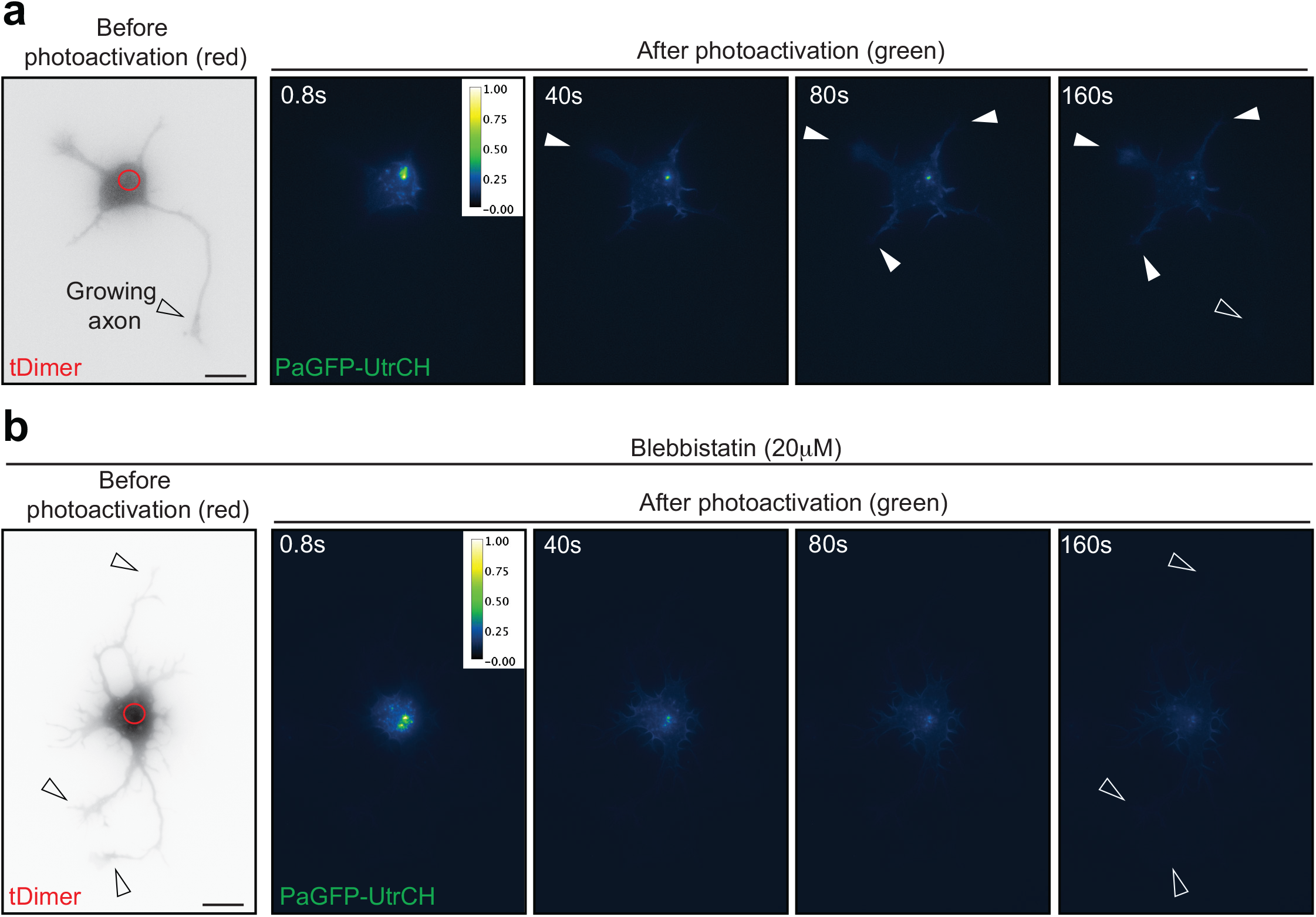
Myosin II inhibition (via 20µM Blebbistatin) affects somatic F-actin delivery. a-b. PaGFP-UtrCH and tDimer co-transfected stage 2 rat hippocampal Control (a) and 20µM Blebbistatin-treated (b) neurons photoactivated in the soma with a 405 nm laser (red circle with a diameter of 5.239 μm).

**Supp Figure 2.**
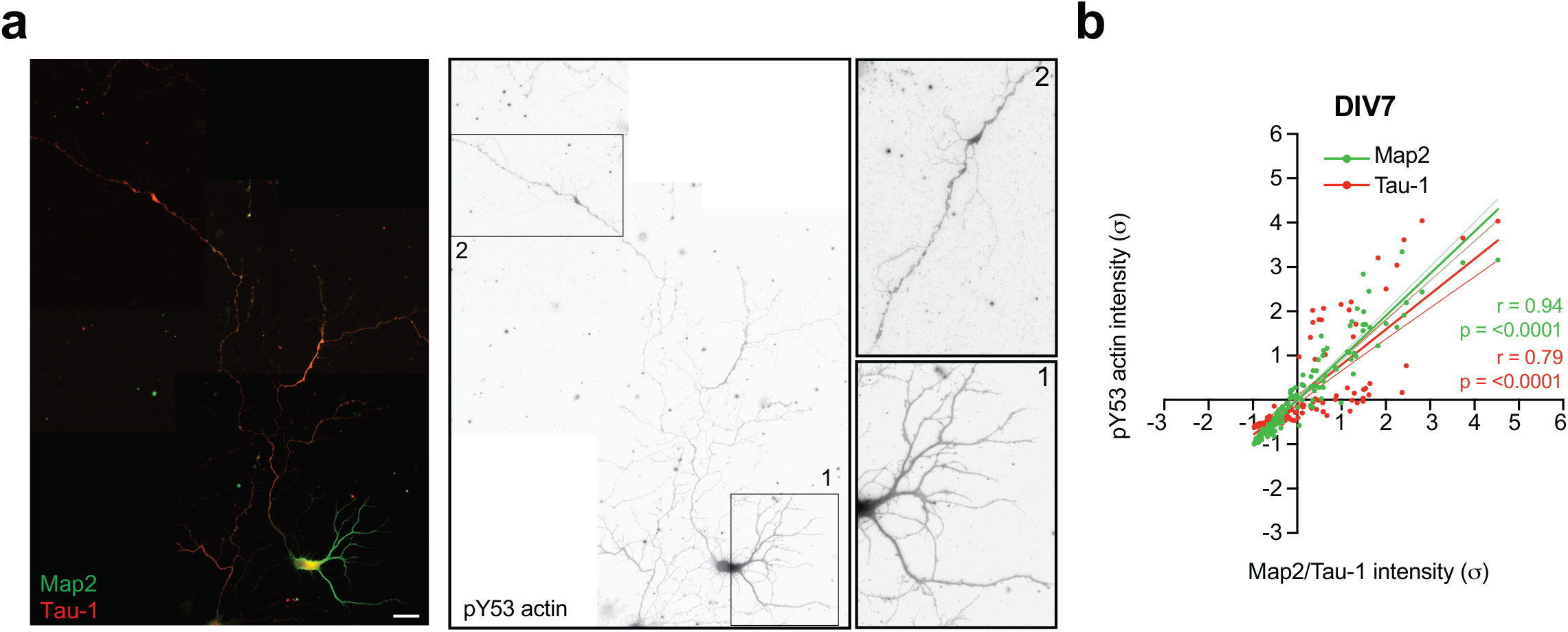
pY53 is eventually enriched in the dendrites of matured neurons (DIV7). a. DIV7 (stage 4) neuron immunostained with pY53 actin, axonal marker Tau-1 and dendritic marker MAP2. b. Linear regression analysis of pY53 actin vs Tau-1 and pY53 actin vs MAP2 intensities from DIV 7 (stage 4) neurons; values were normalized according to a standard score, and axes are represented in units of SD (σ). Linear regression line equation for pY53 actin vs Tau-1 intensities: Y = 0.7952*X - 1.124e-007, pY53 actin vs MAP2 intensities: Y = 0.9498*X - 1.833e-007. Pearson correlation coefficient (r value) and p value of significance are as indicated. Thin lines denote 95% confidence intervals. Data were obtained from n = 21 neurons (153 neurites) from three different cultures.

**Supp Figure 3.**
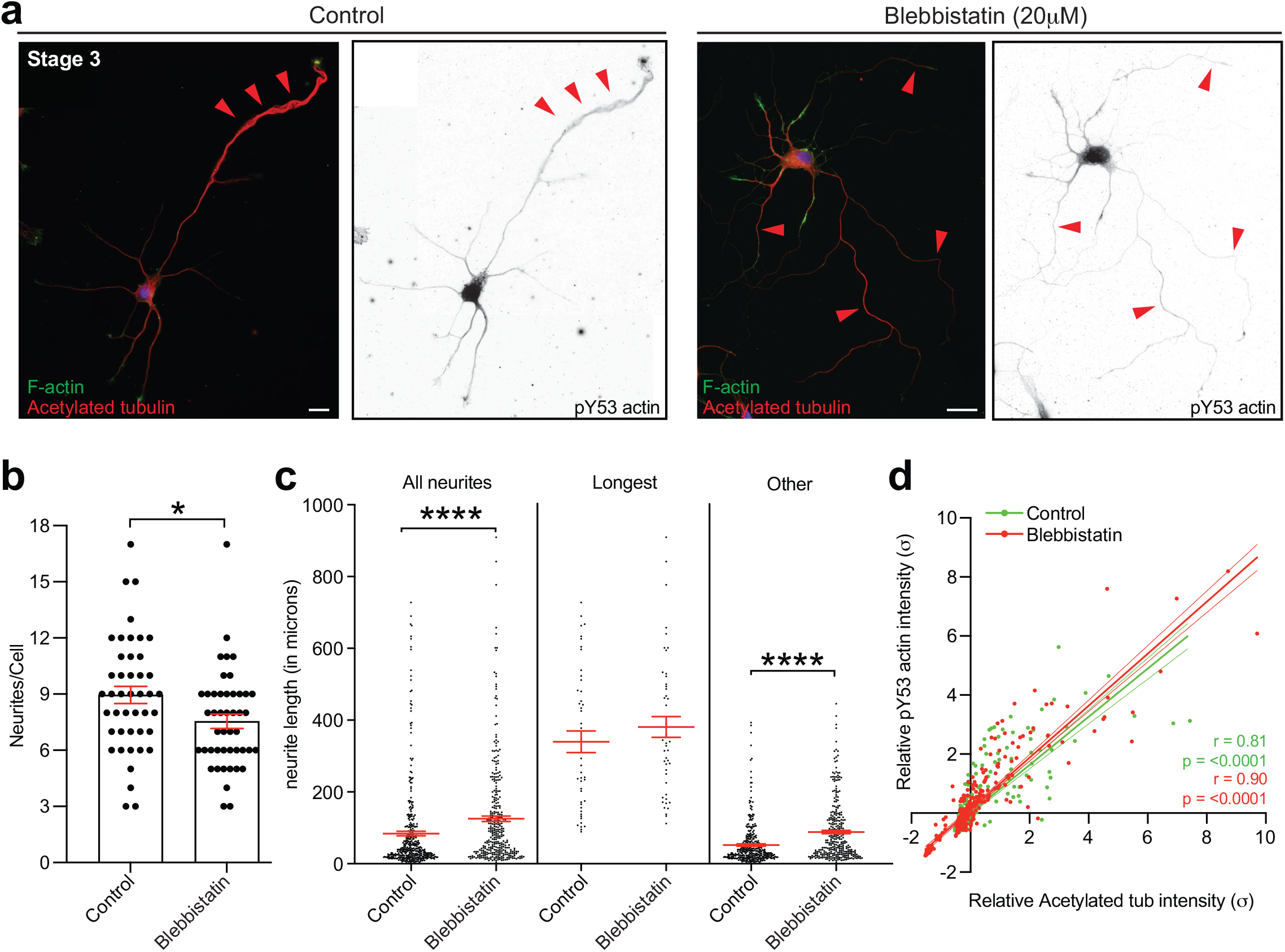
Myosin II inhibition (via 20µM Blebbistatin) results in neurite elongation. a. Control and 20µM Blebbistatin-treated neurons, treated at DIV1 and fixed at DIV3, immunostained with pY53 actin and AcetyTUB and F-actin (labelled by phalloidin-488). b. Quantifications show neurites per cell in Control and 20 µM Blebbistatin-treated neurons, treated at DIV1 and fixed at DIV3. Mean ± SEM neurite number, Control neurons = 8.955 ± 0.4586, Blebbistatin-treated neurons = 7.543 ± 0.3778; * P = 0.0128 by Mann-Whitney non-parametric test. c. Quantifications show neurite length analysis in Control and 20 µM Blebbistatin-treated neurons, treated at DIV1 and fixed at DIV3. Mean ± SEM length values, for all neurites from Control neurons = 83.91 ± 6.275, Blebbistatin-treated neurons = 125.1 ± 7.654; for longest neurites from Control neurons = 339.6 ± 30.14, Blebbistatin-treated neurons = 380.7 ± 28.87; for shorter neurites from Control neurons = 51.98 ± 3.104, Blebbistatin-treated neurons = 88.06 ± 4.518; n.s. = not significant, **** P <0.0001 by Mann-Whitney non-parametric test. d. Linear regression analysis of relative AcetyTUB and pY53 actin intensities in Control and 20 µM Blebbistatin-treated neurons, treated at DIV1 and fixed at DIV3; values were normalized according to a standard score, and axes are represented in units of SD (σ). Linear regression line equation for Control group: Y = 0.8156*X - 1.956e-007, Blebbistatin group: Y = 0.8801*X + 0.1165. Pearson correlation coefficient (r value) and p value of significance are as indicated. Thin lines denote 95% confidence intervals. For data shown in c and d, n = 44 Control cells (383 neurites), n = 46 Blebbistatin-treated cells (332 neurites).

**Supp Figure 4.**
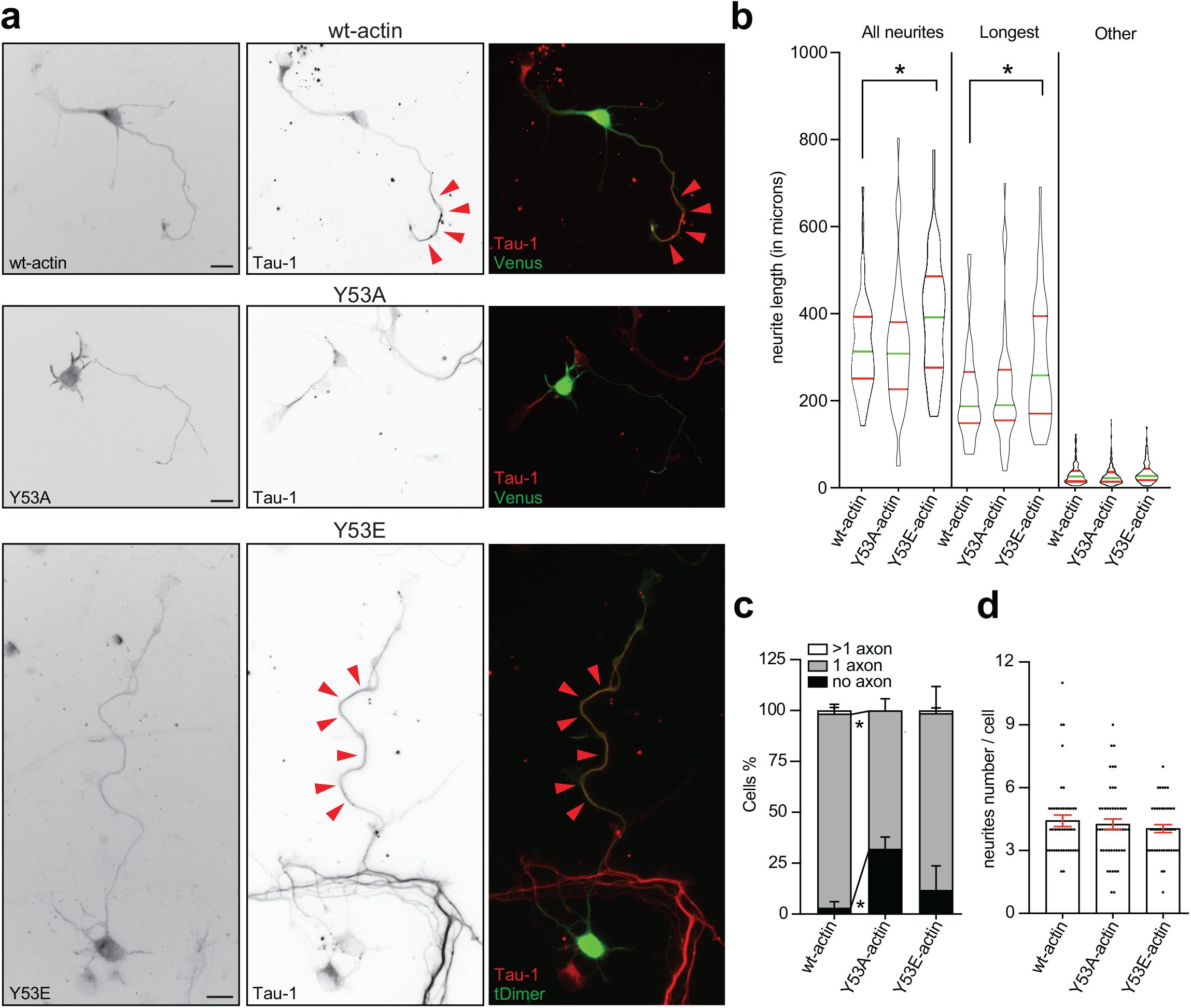
Phosphoactive Y53 actin mutation (Y53E) promotes neurite growth and whereas the phosphodead Y53 actin mutation (Y53A) impairs axon formation. a. Images of mouse cortical neurons co-transfected via IUE at E15 with Venus/tDimer and wt actin or Y53A or Y53E cultured at E17 for 72 h immunostained with Tau-1 to confirm the axonal identity of the neurites, indicated by red arrowheads. b. Quantification of neurite length (µm) for cells expressing wt actin, Y53A or Y53E, as shown in (a). Mean ± SEM values for the length of all neurites in wt actin cells (n = 45) = 326.9 ± 16.69, Y53A cells (n =50) = 324.2± 20.65, Y53E cells (n = 44) = 396.3 ± 21.09. Mean ± SEM values for the length of the longest neurite in wt actin cells = 217.8 ±16.09, Y53A cells = 228.0 ± 19.35, Y53E cells = 290.0 ± 21.69. Mean ± SEM values for length of other neurites in wt actin cells = 31.90 ± 2.018, Y53A cells =28.99 ± 1.879, Y53E cells = 35.19 ± 2.299. P < 0.0001 by one-way ANOVA, post hoc Tukey’s test, * P < 0.05. c. Percentage of neurons expressing wt actin, Y53A or Y53E, as shown in (a), differentiated to have no axon, 1 axon, and >1 axon. Mean ± SEM values for percentage of wt actin (n = 56) cells with no axon = 3.00 ± 3.00, 1 axon = 95.50 ±4.50, more than 1 axon = 1.50 ± 1.50; percentage of Y53A (n = 56) cells with no axon = 32.10 ± 5.80, 1 axon =67.90 ±5.80; percentage of Y53E (n = 55) cells with no axon = 11.80 ± 11.8, 1 axon =86.80 ± 13.20, more than 1 axon =1.30 ± 1.30. a = 0.05 by two-way ANOVA, post hoc Tukey’s test, *P < 0.05. d: Quantification of neurite terminals for cells expressing wt actin, Y53A or Y53E as shown in (a). Mean ± SEM values for neurite terminal per cell in wt actin cells (n = 45) = 4.422 ± 0.2743, Y53A cells (n = 50) = 4.260 ± 0.2488, Y53E cells (n = 44) = 4.045 ± 0.1922. P < 0.0001 by one-way ANOVA test, post hoc Tukey’s test, ns (not significant). Supp Figures 4b, 4c and 4d, data were obtained from cortical cultures of three or more IUE embryos from at least two different mothers.

**Supp Figure 5.**
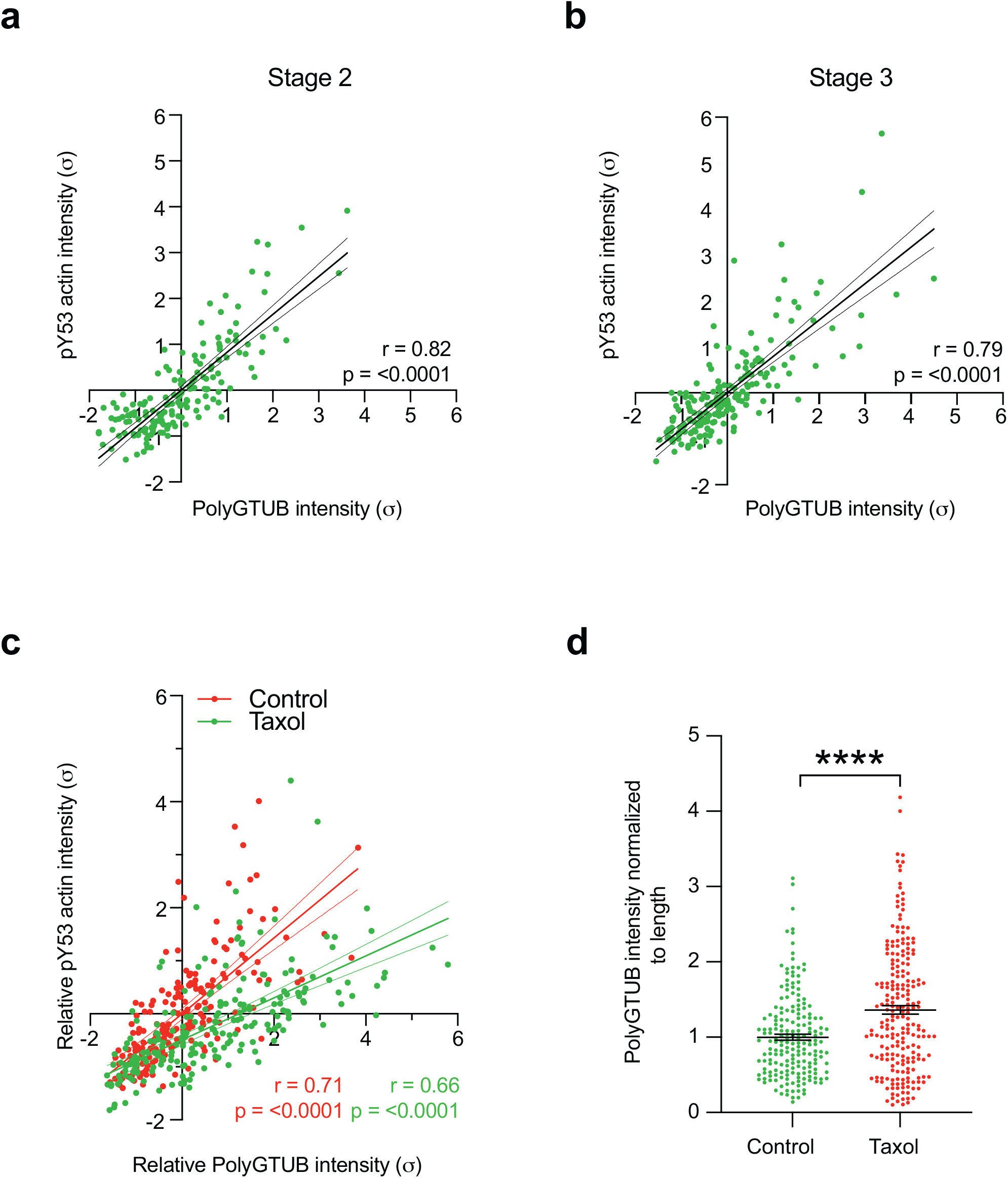
PolyGTUB and pY53 actin distribution correlation in neurite shafts of stage2, stage3 and Taxol-treated neurons. a. Linear regression analysis of PolyGTUB and pY53 actin intensities in stage 2 cells; values were normalized according to a standard score, and axes are represented in units of SD (σ). Linear regression line equation: Y = 0.8620*X + 4.790e-008. Pearson correlation coefficient (r value) and p value of significance are as indicated. Thin lines denote 95% confidence intervals. Data were obtained from 167 neurites of 38 neurons from two different cultures. b. Linear regression analysis of PolyGTUB and pY53 actin intensities in stage 3 cells; values were normalized according to a standard score, and axes are represented in units of SD (σ). Linear regression line equation Y = 0.7925*X + 1.284e-007. Pearson correlation coefficient (r value) and p value of significance are as indicated. Thin lines denote 95% confidence intervals. Data were obtained from 194 neurites of 22 neurons from three different cultures. c. Linear regression analysis of PolyGTUB and pY53 actin intensities in Control and Taxol (neurons incubated with 5nM Taxol 24h after plating for 48h) groups; values were normalized according to a standard score, and axes are represented in units of SD (σ). Linear regression line equations for pY53 actin intensities from Control group: Y = 0.7151*X - 5.742e-007; from Taxol group: Y = 0.3959*X - 0.4918. Pearson correlation coefficient (r value) and p value of significance are as indicated. Thin lines denote 95% confidence intervals. d. Quantifications from Control and Taxol (neurons incubated with 5nM Taxol 24h after plating for 48h) groups. Quantification of relative PolyGTUB intensity values normalized to length from the neurite shafts. Mean ± SEM values from Control neurons = 1.000 ± 0.03952 and 48h Taxol-treated neurons = 1.361 ± 0.05624. **** P <0.0001 by Mann-Whitney non-parametric test. Supp Figure 5c and 5d, data were obtained from 194 neurites of 22 Control neurons and 220 neurites of 27 Taxol-treated (48h) neurons from three different cultures.

**Supp Figure 6.**
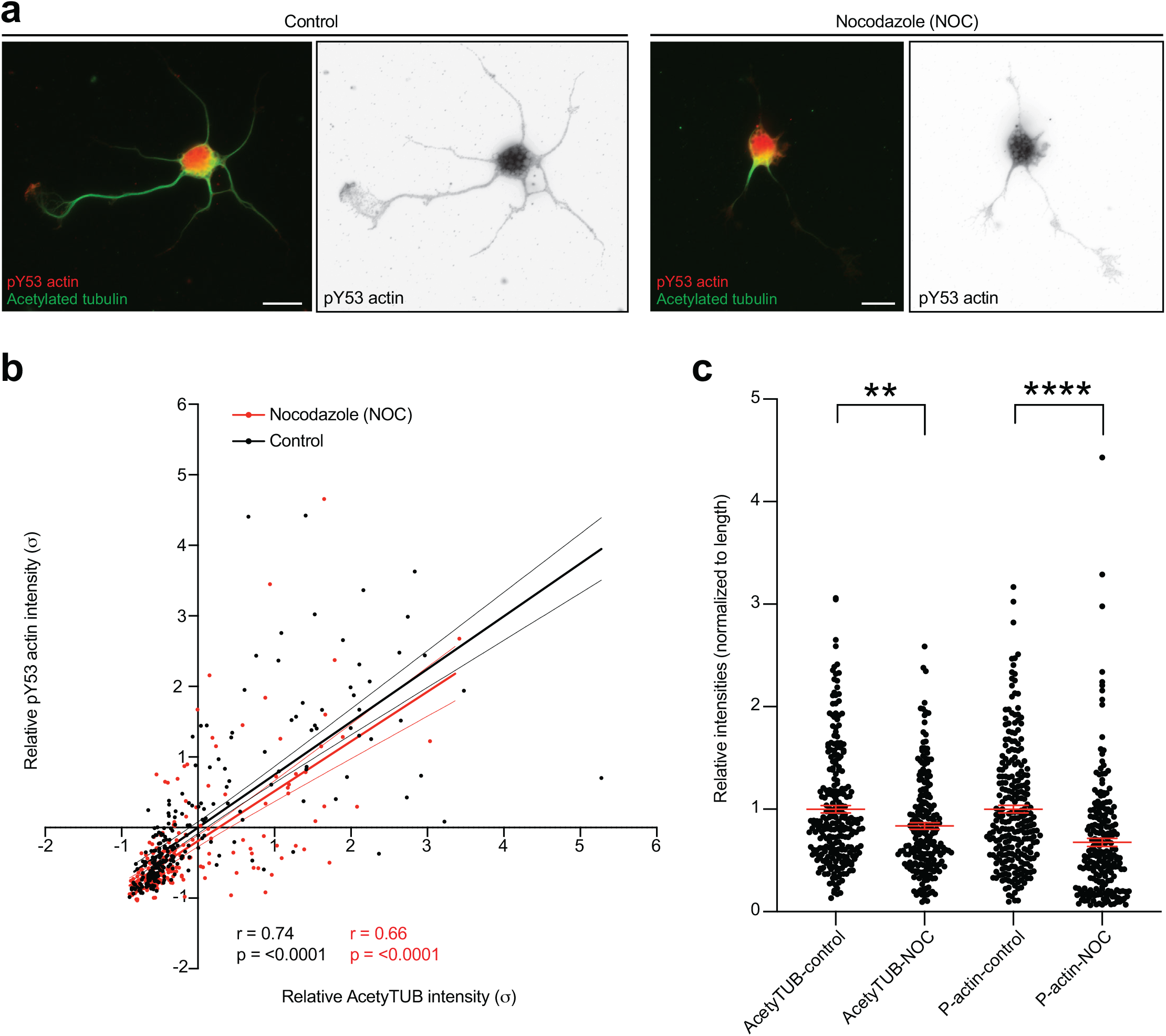
Nocodazole-induced MT destabilization decreases pY53 actin levels in DIV1 neurons. a. Control and 6µM Nocodazole-treated neurons, treated at DIV1 for 30 min, immunostained with pY53 actin and AcetyTUB. b. Linear regression analysis of relative AcetyTUB and pY53 actin intensities in Control and 6 µM Nocodazole-treated neurons; values were normalized according to a standard score, and axes are represented in units of SD (σ). Linear regression line equation for the Control group: Y = 0.7486*X - 8.957e-008, Nocodazole group: Y = 0.7051*X - 0.1897. Pearson correlation coefficient (r value) and p value of significance are as indicated. Thin lines denote 95% confidence intervals. c. Quantification of AcetyTUB and pY53 actin intensity values normalized to length from neurites (per each cell). Mean ± SEM values for normalized AcetyTUB intensity in the Control group = 1.000 ± 0.03592 and Nocodazole-treated group= 0.8360 ± 0.03182. ** P = 0.0025 by Mann-Whitney non-parametric test. Mean ± SEM values for normalized pY53 actin intensity in the Control group = 1.000 ± 0.03695 and Nocodazole-treated group = 0.6758 ± 0.03851. **** P <0.0001 by Mann-Whitney non-parametric test. Supp Figures 6b and 6c, data were obtained from 61 Control cells (252 neurites), n = 56 Nocodazole-treated cells (219 neurites) from three different cultures.

**Movie EV1. Photoactivation in the soma of stage 2 neuron expressing PaGFP-UtrCH and tDimer.**

Imaging was performed on a FRAP imaging system based on a Nikon Ti-E equipped with a Nikon CFI Apo TIRF 100x, 1.49 NA oil objective. 405 nm laser illumination is performed in a circular region with a dimeter of 5.239 µm to achieve photoactivation in the soma. Duration of time-lapse imaging: 160 sec; 2.4 sec before and 157.6 sec after photoactivation. Interval between the frames is 0.8 sec.

**Movie EV2. Photoactivation in the soma of stage 3 neuron expressing PaGFP-UtrCH and tDimer.**

Imaging was performed on a FRAP imaging system based on a Nikon Ti-E equipped with a Nikon CFI Apo TIRF 100x, 1.49 NA oil objective. 405 nm laser illumination is performed in a circular region with a dimeter of 5.239 µm to achieve photoactivation in the soma. Duration of time-lapse imaging: 160 sec; 2.4 sec before and 157.6 sec after photoactivation. Interval between the frames is 0.8 sec.

**Movie EV3. Photoactivation in the soma of stage 4 neuron expressing PaGFP-UtrCH and tDimer.**

Imaging was performed on a FRAP imaging system based on a Nikon Ti-E equipped with a Nikon CFI Apo TIRF 100x, 1.49 NA oil objective. 405 nm laser illumination is performed in a circular region with a dimeter of 5.239 µm to achieve photoactivation in the soma. Duration of time-lapse imaging: 160 sec; 2.4 sec before and 157.6 sec after photoactivation. Interval between the frames is 0.8 sec.

**Movie EV4. Photoactivation in the soma of stage 2 neuron expressing PaGFP-Myosin IIA and mMaroon1.**

Imaging was performed on a FRAP imaging system based on a Nikon Ti-E equipped with a Nikon CFI Apo TIRF 100x, 1.49 NA oil objective. 405 nm laser illumination is performed in a circular region with a dimeter of 5.239 µm to achieve photoactivation in the growth cone. Duration of time-lapse imaging: 120 sec; 2.4 sec before and 117.6 sec after photoactivation. Interval between the frames is 0.8 sec.

**Movie EV5. Photoactivation in the soma of the neuron expressing PaGFP-UtrCH and tDimer treated with 20µM Blebbistatin.**

Imaging was performed on a FRAP imaging system based on a Nikon Ti-E equipped with a Nikon CFI Apo TIRF 100x, 1.49 NA oil objective. 405 nm laser illumination is performed in a circular region with a dimeter of 5.239 µm to achieve photoactivation in the growth cone. Duration of time-lapse imaging: 160 sec; 2.4 sec before and 157.6 sec after photoactivation. Interval between the frames is 0.8 sec.

**Movie EV6. Photoactivation in the soma of stage 2 neuron expressing PaGFP-UtrCH and mMaroon1.**

Imaging was performed on a FRAP imaging system based on a Nikon a Nikon EclipseTi2 inverted microscope quipped with a PLAN APO 60X oil (NA 1.42) objective. 405 nm laser illumination is performed in a circular region with a dimeter of 5.239 µm to achieve photoactivation in the growth cone. Duration of time-lapse imaging: 120 sec; 2.4 sec before and 117.6 sec after photoactivation. Interval between the frames is 0.8 sec.

**Movie EV7. Photoactivation in the soma of stage 2 neuron expressing PaGFP-UtrCH and Kif5c-Tdtomato.**

Imaging was performed on a FRAP imaging system based on a Nikon EclipseTi2 inverted microscope quipped with a PLAN APO 60X oil (NA 1.42) objective. 405 nm laser illumination is performed in a circular region with a dimeter of 5.215 µm to achieve photoactivation in the neurites. Duration of time-lapse imaging: 120 sec; 2.4 sec before and 117.6 sec after photoactivation. Interval between the frames is 0.8 sec.

